# Pre-movement sensor-level EEG activity recorded with a portable three-channel system is associated with futsal free-kick outcomes

**DOI:** 10.64898/2025.12.11.691239

**Authors:** Sadao Hirose, Junichi Ushiba, Seitaro Iwama

## Abstract

Brain states immediately before movement may shape skilled athletic performance, but translating electroencephalography (EEG)-based monitoring to sport has been limited by dense montages and laboratory-constrained tasks. We asked whether a portable three-channel EEG system can detect outcome-related preparatory activity during a realistic futsal free-kick task. EEG from C3, Cz, and C4 during the 2 s before each kick was used to decode successful versus unsuccessful trials with channel-wise linear support vector machines (SVMs), and the same approach was tested for transfer across recording days. To relate decoding to interpretable physiology, alpha- and beta-band power were also compared between outcomes. Successful and failed kicks were decoded significantly above label-shuffled null distributions within recording days across all channels, showing that low-density EEG can capture sensor-level information linked to subsequent performance. However, decoding did not generalize reliably across days, indicating that these neural signatures were session specific. Spectral analysis showed selectively increased C3 beta-band power before successful kicks, whereas alpha-band power showed no significant outcome-related differences. These findings support portable EEG as a practical route toward monitoring preparatory brain states in sport, while suggesting that real-world use will require short day-specific calibration or adaptive decoding.

## Introduction

The primary objective of the sports science field is to enhance the competitive performance of athletes[1]. While traditional approaches have focused on the development of physical and technical skills, current work has shifted toward the continuous monitoring and active regulation of the athletes’ internal physiological states, with the goal of optimizing training quality and, ultimately, elevating competitive outcomes. Specifically, the quantification of fatigue using heart-rate variability, recovery kinetics, and blood-lactate concentrations allows practitioners to titrate external training loads, while immersive virtual-reality platforms are being employed to refine the athletes’ perceptual-cognitive and decision-making skills during sport-specific scenarios[2,3]. Due to rapid advances in wearable sensors, data analytics, and extended-reality hardware, these monitoring and simulation technologies have become ubiquitous, particularly in elite and professional settings, where their integration in daily practice is almost standard[4–6].

Recent attention has turned to the potential benefits of systematically monitoring and managing brain states to enhance performance. Neural regulation may therefore be a promising approach for the optimization of athletic performance[7]. One line of research emphasizing the influence of brain activity on sports performance focuses on controlling pre-movement brain activity. Previous findings indicate that self-regulation training can improve motor neural activities during the pre-movement period[8–11]. For example, Iwama et al.[9] demonstrated that short-term neurofeedback training based on scalp electroencephalogram (EEG) from the sensorimotor cortex enhanced its excitability and improved fine motor control in a touch-typing task, which highlighted the practical benefits of modulating preparatory neural activity. Complementary studies have attempted to predict athletic performance by decoding the preparatory neural state. These investigations demonstrated that the likelihood of success was enhanced when movement was initiated at an “optimal” configuration for cortical activity[12–14]. For example, Meinel et al.[13] showed that initiating movement during specific oscillatory brain states, identified via EEG-based closed-loop gating, significantly improved motor performance in chronic stroke patients. Collectively, this body of evidence suggests that actively shaping, or strategically timing action to coincide with, an optimal preparatory neural state immediately before movement can yield measurable gains in motor performance.

Despite this progress, there remain practical barriers to applying EEG-based performance prediction in real-world sports settings. Notably, several research investigations have employed high-density EEG systems, which make it challenging to capture brain activity during actual physical movements. High-density EEG systems are often time-consuming to set up and can constrain natural movement during use, rendering them impractical for deployment in training sessions or competitive environments. Additionally, validation of EEG-based performance analysis has mostly been conducted under organized laboratory conditions, limiting its direct applicability within real-world sports environments. Since there are substantial differences in the athletes’ psychological and physiological states between laboratory settings and actual competitive environments, the external validity of findings obtained under controlled conditions remains limited.

In this study, we address these challenges by examining a soccer-specific skill—kicking a ball to a fixed target that closely approximates real competitive actions. Using a portable 3-channel EEG system, we recorded pre-movement EEG activity and developed classification models to predict whether a given kick would succeed or fail on a single-trial basis. Thus, this study tested (i) the feasibility of performance prediction with a lightweight EEG device and (ii) the applicability of this approach under conditions approximating real-world sports scenarios. By incorporating a practical workflow from data acquisition to prediction, our study lays the foundation for future training protocols that could actively shape the brain states of athletes for optimal performance enhancement.

## Methods

### Ethics statement

The study protocol was approved by the Ethics Committee of the Faculty of Science and Technology at Keio University (approval numbers: 2023-141, 2024-011) and conducted in accordance with the Declaration of Helsinki. Written informed consent was obtained from each participant before the experiment. The participant shown in Fig 1 is marked in blue and has given written informed consent for publication of the image. The individual visible in the background is an author of this article and provided written informed consent for publication of the image.

**Fig 1.**
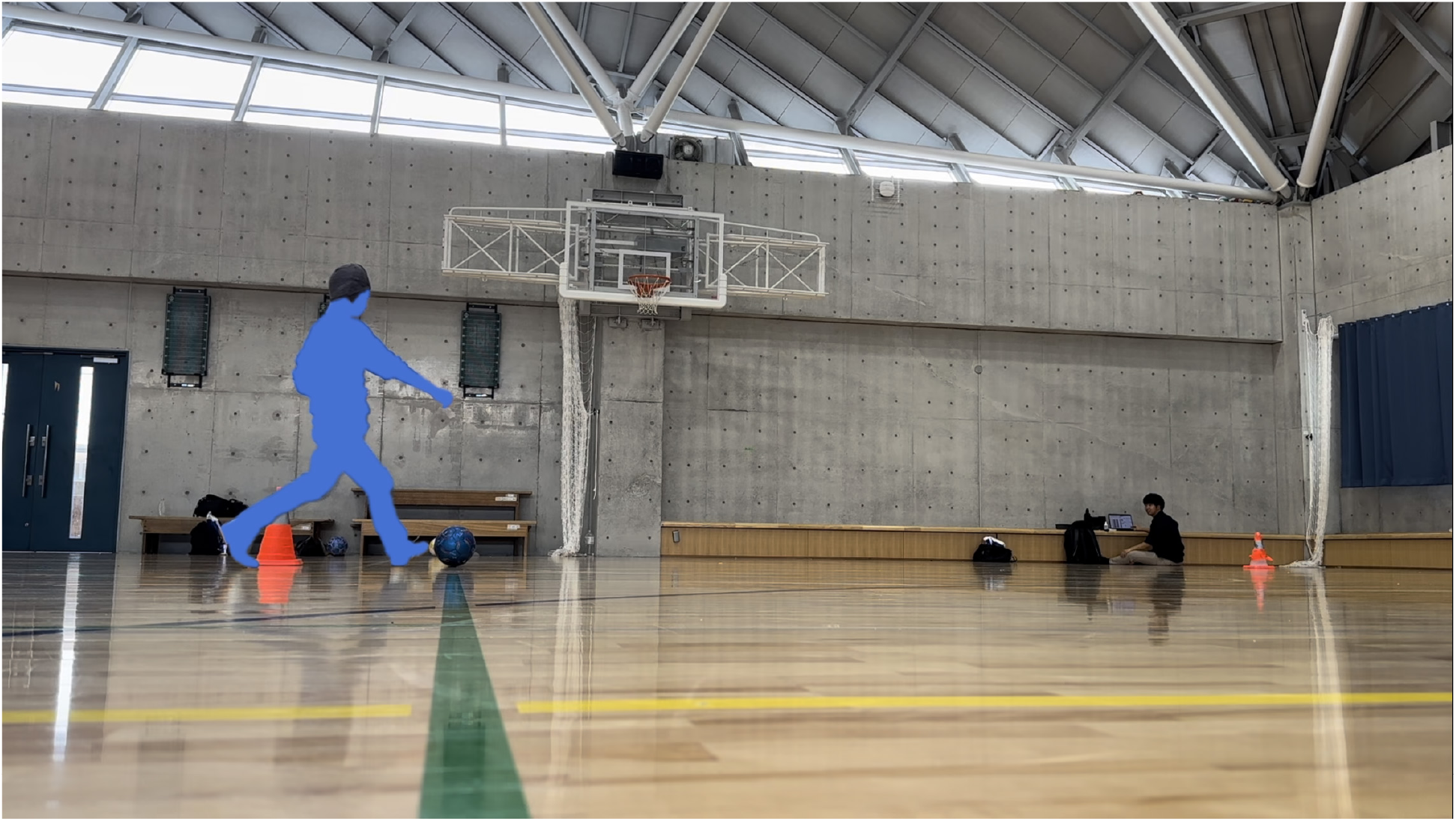
Experimental setup for the kicking task. The right-footed participant marked in blue is equipped with the 3-channel headphone-type EEG system (C3, Cz, C4) and prepares to kick a FIFA size 4 futsal ball 10.4 m toward a square-pyramid cone inside the university gymnasium. The individual visible in the background is an author of this article. Photograph by Sadao Hirose, licensed under CC BY 4.0.

### Participants

In this study, the participants were 15 right-footed males aged 19 to 21 years (19.3±0.6) who had been playing soccer for more than 5 years. Data measurements were taken at the University of Tokyo gymnasium. Participants were recruited between 15 March 2024 and 15 May 2024.

### Data acquisition

The experiment was designed to retain key field-relevant features of futsal kicking while allowing reliable recording of pre-movement EEG. We used a custom-made headphone-shaped 3-channel EEG setup. The EEG amplifier was derived from a commercially available EEG system (AE-120A, Nihon Kohden, Japan). EEG signals from the C3, C4, and Cz channels and head acceleration were simultaneously recorded at a sampling rate of 200 Hz. The impedance of each electrode was kept below 50 kΩ via manual adjustment. Participants underwent two experimental sessions on separate days and completed 50 trials per day, yielding 100 trials per participant and 1,500 trials in total. As specified in futsal laws, participants kicked a Fédération Internationale de Football Association (FIFA)-regulation size 4 futsal ball (circumference: 62–64 cm; weight: 390–430 g; bounce: 55–65 cm) toward a cone shaped like a square pyramid (40 cm in height, 24 cm in width) that was placed at a distance of 10.4 m. Although the gymnasium was not solely reserved for the experiment and a few other individuals were engaged in exercise activities, the experimental setup was arranged to ensure that they remained outside the participant’s field of vision. EEG was recorded from 5 s before the kicking motion began until the kick was completed. To reduce movement-related contamination during the pre-movement EEG period, preparatory movement was limited to a single step. Although this excluded long run-ups, the resulting short-approach kick was consistent with many passes in actual soccer and futsal matches that are performed with a single preparatory step rather than a multi-step run-up. Approximately 40 s were required between kicks to retrieve the ball and set it up again. It was also recorded whether the ball hit the cone or not. Fig 1 shows the setup during the kicking trials.

### EEG signal preprocessing

The data preprocessing flow is illustrated in Fig 2. Based on the acquired data, segments immediately before the ball was kicked were extracted, filtered, and only the appropriate data were used for analysis. The search for movement onset began after the first 400 samples (2 s at the 200 Hz sampling rate) to ensure that the 2 s pre-movement segment required for subsequent analyses was available. Movement onset was identified as the first non-zero sample at which the absolute deviation of the head acceleration signal from the mean of the first 100 non-zero acceleration values exceeded 2 m/s^2^. Only samples preceding the detected onset were retained. Trials were included only when electrode impedance remained below 50 kΩ and no clear movement artifacts were observed in the pre-movement waveform. The acceleration signal was used solely to identify movement onset and was not included as an input to the classifier. The signal was first processed using a 50 Hz notch filter to remove power line interference. Subsequently, a third-order Butterworth bandpass filter (3–70 Hz) was applied to extract the desired frequency components. Both filters were applied as forward-only causal filters. The last 2 s of frames were extracted. A representative 10 s EEG waveform and the extracted 2 s pre-movement segment are shown in Fig 3a and Fig 3b, respectively. Power spectral densities (PSDs) were estimated for each trial and channel, and the resulting spectra were parameterized using Fitting Oscillations & One-Over-F (FOOOF; version 1.1.0)[15] to assess spectral quality and aperiodic components. One trial from one participant was excluded from subsequent EEG analyses because its spectral fit yielded a negative aperiodic exponent, leaving 1,499 trials for analysis.

**Fig 2.**
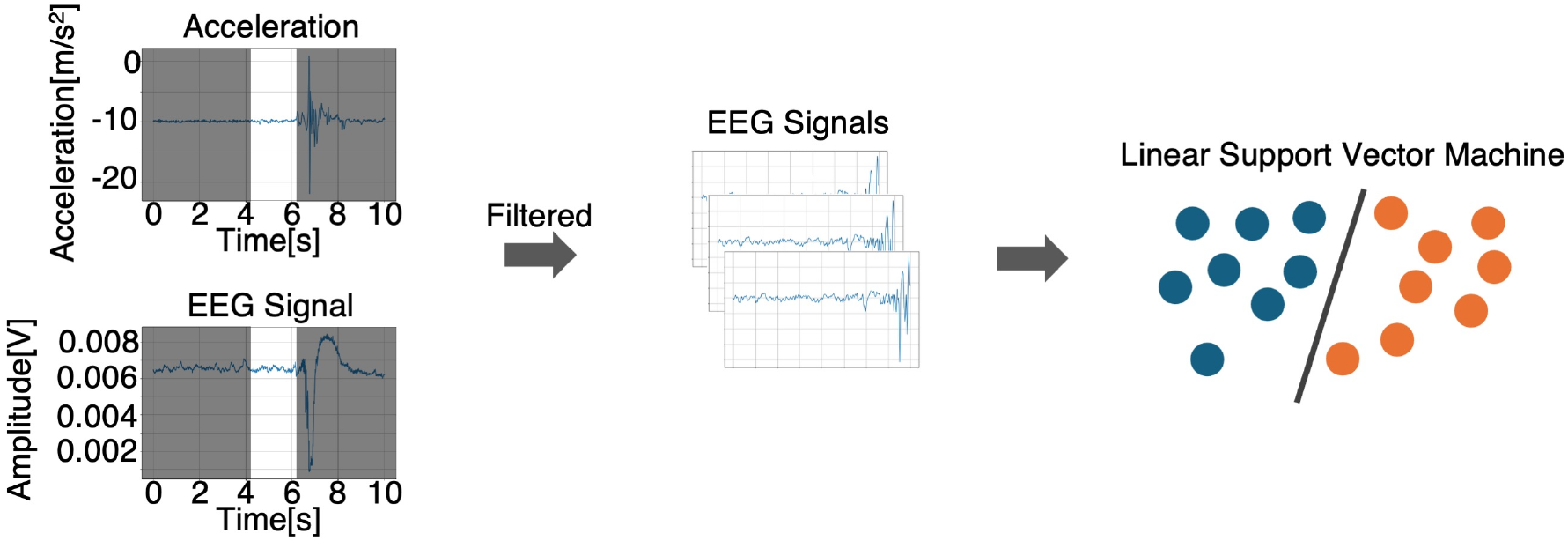
EEG preprocessing workflow. Step-by-step illustration of the data preprocessing procedure, starting from the extraction of relevant segments from the raw data to the final feature generation.

**Fig 3.**
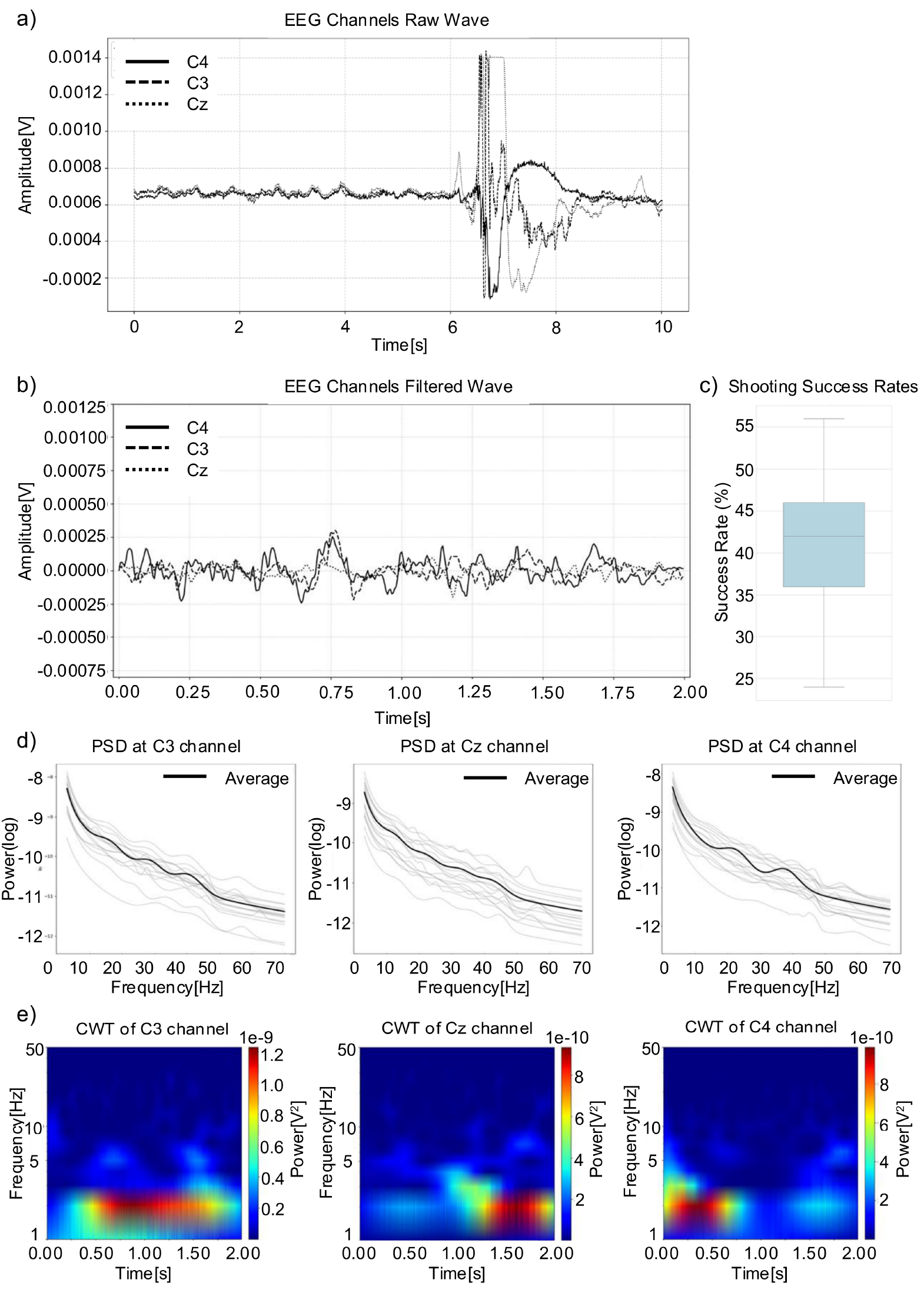
Representative EEG waveforms, behavioral performance, spectral characteristics, and time-frequency representations. a) Raw EEG waveform over a 10 s period, including both before and after the kicking movement. b) Representative 2 s segment from one trial after filtering, displayed for all three channels (C3, Cz, C4), with time 2.00 indicating the movement onset. c) Boxplot of shooting success rates across participant-day observations. d) PSD of pre-movement EEG. Thin colored curves are individual participants; thick black curve is the across-participant mean. e) Grand-average Morlet wavelet power (1–50 Hz) across all trials, computed using the Continuous Wavelet Transform (CWT).

### Linear support vector machine (SVM)-based decoding of kick outcomes

The analytical approach was revised from the original version of this preprint (doi: 10.64898/2025.12.11.691239), which used a transformer-based encoder with a multilayer perceptron decoder. Given the limited sample of 15 participants assessed on two days, we adopted a linear SVM to test outcome-related pre-movement EEG information more directly and interpretably at the sensor level. Label-shuffle permutation testing and Holm correction were used to prioritize conservative statistical inference over predictive performance.

To determine whether kick outcomes could be decoded from pre-movement EEG signals using a linear model, we constructed a linear support vector machine (linear SVM) decoder. EEG signals were analyzed separately for each electrode. Each trial was represented as a 400-point one-dimensional waveform corresponding to the 2 s pre-movement period and was used as input to the classifier.

For each participant, recording day, and EEG channel, classification performance was evaluated using four-fold stratified cross-validation. Within each fold, waveform amplitudes were standardized using parameters estimated only from the training split, and the same transformation was applied to the corresponding validation split. A class-balanced linear SVM was used for classification to account for class imbalance between successful and unsuccessful trials. The regularization parameter (C) was optimized separately for each participant, recording day, and channel by grid search over ({0.01, 0.1, 1, 10, 100}). For each value of (C), the model was trained for up to 10,000 iterations, with optimization terminated earlier if the solver convergence criterion was satisfied. The value of (C) that yielded the highest mean validation macro-F1 score across the four folds was selected. When multiple values produced the same macro-F1 score, the model with the lower hinge loss was selected.

Model performance was primarily evaluated using the mean area under the receiver operating characteristic curve (ROC AUC) across the four validation folds, because AUC provides a threshold-independent measure of discriminability. To assess whether the observed decoding performance exceeded chance level, we performed a label-shuffle permutation test separately for each combination of participant, recording day, and EEG channel. In each permutation, trial labels were randomly shuffled while preserving the numbers of successful and unsuccessful trials. The regularization parameter (C) selected in the within-day SVM analysis for the corresponding participant, recording day, and channel was reused, and the same four-fold cross-validation procedure was repeated using the shuffled labels. A total of 199 permutations were performed for each combination. Empirical upper-tail p values were computed by comparing the observed AUC with the null distribution obtained from the shuffled-label analyses. Channel-level statistical summaries were then computed by comparing the observed across-participant mean AUC with the corresponding permutation-based null distribution, with Holm correction applied across channels.

### Across-day generalization analysis

To evaluate whether the linear SVM decoder generalized across recording days, we performed an across-day classification analysis. For each participant and EEG channel, two directional evaluations were conducted: training on Day 1 and testing on Day 2, and training on Day 2 and testing on Day 1. In all analyses, data from the evaluation day were not used for model selection, hyperparameter tuning, preprocessing-parameter estimation, or model training.

For the linear SVM decoder, data from the training day were divided into four stratified folds. The regularization parameter (C) selected in the within-day SVM analysis for the corresponding participant, training day, and channel was reused. In each fold, waveform amplitudes were standardized using parameters estimated only from the training split, and a class-balanced linear SVM was fitted using the standardized training data. The trained model was then applied to all trials from the evaluation day to obtain decision scores. For each evaluation-day trial, decision scores were averaged across the four fold-specific models, and the final across-day ROC AUC was computed from these averaged decision scores. To assess chance-level performance, the evaluation-day labels were randomly shuffled 199 times while preserving the numbers of successful and unsuccessful trials. The observed across-day AUC was then compared with the null distribution obtained from the shuffled-label analyses.

Across-day performance was primarily evaluated using ROC AUC, because AUC provides a threshold-independent measure of discriminability. Empirical upper-tail p values were computed by comparing the observed across-day AUC with the corresponding shuffled-label null distribution. Channel-level summaries were computed by averaging performance across participants and evaluation directions, and Holm correction was applied across channels.

### Examination of alpha- and beta-band oscillations during the pre-movement period

To examine whether oscillatory activity during the pre-movement period differed between successful and unsuccessful kicks, we analyzed alpha- and beta-band power separately for each EEG channel. The alpha and beta bands were defined as 8–15 Hz and 15–30 Hz, respectively. For each participant and channel, trials from both recording days were pooled and divided into successful and unsuccessful trials according to the behavioral outcome of each kick.

For each participant, channel, and outcome condition, EEG waveforms were first averaged across trials to obtain representative pre-movement waveforms for successful and unsuccessful kicks. A fast Fourier transform was then applied to each averaged waveform, and the power spectrum was computed as the squared magnitude of the Fourier coefficients. Band power was calculated by summing spectral power within each frequency band and multiplying by 2/N^2^ for one-sided normalization (N = 400 samples). This procedure yielded one success value and one failure value for each participant, channel, and frequency band, thereby preserving the participant as the unit of statistical analysis.

For each channel and frequency band, band-power values for successful and unsuccessful kicks were compared across participants using paired statistical tests. The normality of the paired differences was assessed using the Shapiro–Wilk test. When the paired differences were normally distributed, a paired t-test was performed. Otherwise, a Wilcoxon signed-rank test was used. Because alpha- and beta-band activities were treated as separate frequency-specific hypothesis families, multiple comparisons were corrected separately within each band. Specifically, Holm correction was applied across the three channel-wise comparisons within the alpha band and across the three channel-wise comparisons within the beta band.

All analyses were performed in Python 3.8.16 using SciPy version 1.10.1 for signal processing and statistical tests, and scikit-learn version 1.3.2 for the SVM analyses[16,17].

## Results

### Behavioral Performance and EEG Data Quality Assessment

Following spectral-quality screening, 1,499 of the 1,500 recorded trials were retained for EEG analysis. The proportion of successful trials for each participant across days is presented in Fig 3c.

The results of the EEG data quality check are presented in Fig 3. Participant-level PSDs and the corresponding across-participant averages are shown for each channel in Fig 3d. Power spectra were modeled using the FOOOF algorithm. Fits were applied to the 3–70 Hz range using the following settings: peak_width_limits = [2, 8] Hz, max_n_peaks = 6, and min_peak_height = 0.15 (relative to the aperiodic background). Fit quality was uniformly high across all electrodes. For the C4 channel, FOOOF achieved a median R^2^ of 0.980 (interquartile range (IQR) = 0.006) and a median mean absolute error (MAE; log10-power space) of 0.077 (IQR = 0.008). Comparable performances were obtained for the C3 channel (median R^2^ = 0.979, IQR = 0.014; median MAE = 0.077, IQR = 0.022) and Cz channel (median R^2^ = 0.986, IQR = 0.010; median MAE = 0.069, IQR = 0.019). Additionally, for each electrode, trials were averaged and submitted to a continuous Morlet wavelet transform using 50 scales corresponding to equally spaced frequencies from 1 to 50 Hz. In Fig 3e, power (|CWT|^2^) was visualized as a time-frequency map with a logarithmic frequency axis to confirm that canonical EEG rhythms were preserved.

### Successful and Failed Trials Are Decodable Within Days but Show Limited Cross-Day Generalization

To examine whether single-channel EEG activity contained information about kick outcome, we evaluated the channel-wise linear SVM classifier. The classification results are summarized in Fig 4. In the within-day analysis, the linear SVM achieved AUC values significantly higher than the label-shuffled null distribution for all three channels (Fig 4a; C3: observed AUC = 0.561, null mean = 0.500, Holm-adjusted p = 0.015; Cz: observed AUC = 0.556, null mean = 0.502, Holm-adjusted p = 0.015; C4: observed AUC = 0.577, null mean = 0.499, Holm-adjusted p = 0.015).

**Fig 4.**
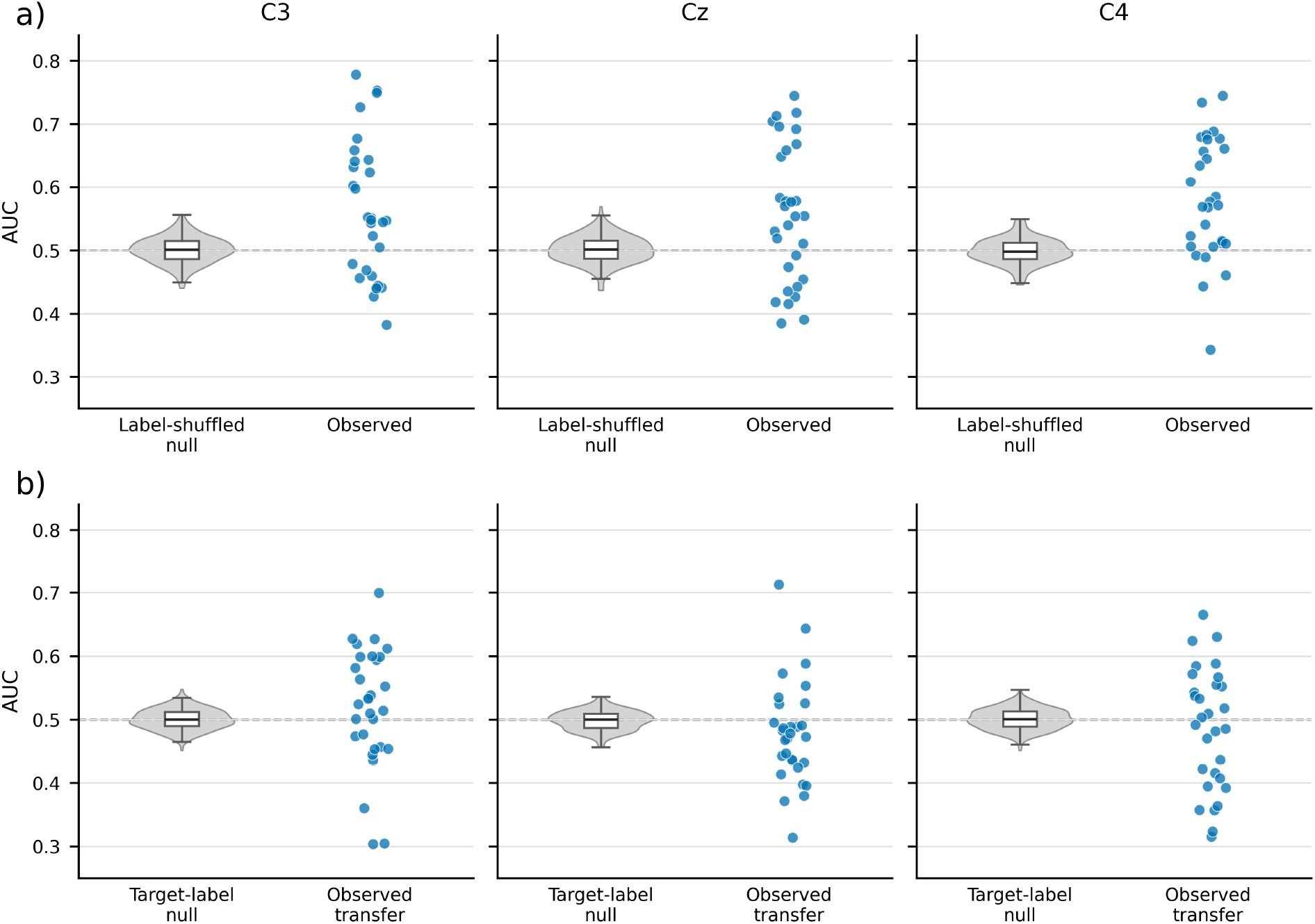
Classification AUC across within-day and cross-day analyses using the linear SVM classifier. Panels a and b show SVM within-day and SVM cross-day results, respectively. Columns show C3, Cz, and C4. For the within-day analysis, each blue point represents the mean cross-validated AUC for one participant on one recording day. For the cross-day analysis, each blue point represents the AUC for one participant in one evaluation direction, either Day 1 to Day 2 or Day 2 to Day 1. Gray violins show the channel-level permutation null distributions obtained from 199 permutations after averaging AUCs across the corresponding participant-day units for the within-day analysis or participant-direction units for the cross-day analysis. Overlaid boxplots summarize the null distributions. The dashed horizontal line marks AUC = 0.5. a) SVM within-day: C3 observed mean AUC = 0.561, null mean = 0.500; Cz observed mean AUC = 0.556, null mean = 0.502; C4 observed mean AUC = 0.577, null mean = 0.499. b) SVM cross-day: C3 observed mean AUC = 0.520, null mean = 0.500; Cz observed mean AUC = 0.479, null mean = 0.499; C4 observed mean AUC = 0.487, null mean = 0.501.1

However, this decoding performance did not generalize reliably across days. When the model trained on one day was evaluated on the other day, SVM AUCs did not exceed the target-label permutation null distribution after Holm correction (Fig 4b; C3: observed AUC = 0.520, null mean = 0.500, Holm-adjusted p = 0.405; Cz: observed AUC = 0.479, null mean = 0.499, Holm-adjusted p = 1.000; C4: observed AUC = 0.487, null mean = 0.501, Holm-adjusted p = 1.000). These results indicate that successful and failed kicks were decodable from single-channel EEG within the same day, whereas the discriminative patterns were not stably transferable across days.

### Successful Kicks Were Preceded by Selectively Increased C3 Beta-Band Power

Band-power analysis revealed a frequency- and channel-specific difference between successful and unsuccessful kicks, as summarized in Fig 5. In the beta-band analysis, C3 beta-band power was significantly greater before successful than unsuccessful kicks after Holm correction (Fig 5b; success: 48.9 µV^2^; failure: 32.7 µV^2^; paired t-test, t(14) = 2.81, two-sided p = 0.0139, Holm-adjusted p = 0.0418). In contrast, the remaining beta-band comparisons were not significant after correction (Cz: Holm-adjusted p = 0.375; C4: Holm-adjusted p = 0.524). In the alpha-band analysis, no channel showed a significant difference after Holm correction (C3: Holm-adjusted p = 0.381; Cz: Holm-adjusted p = 0.191; C4: Holm-adjusted p = 0.528). These results indicate that successful kicks were associated with a selective increase in pre-movement beta-band power over C3, rather than a broad increase across all central channels or across both frequency bands.

**Fig 5.**
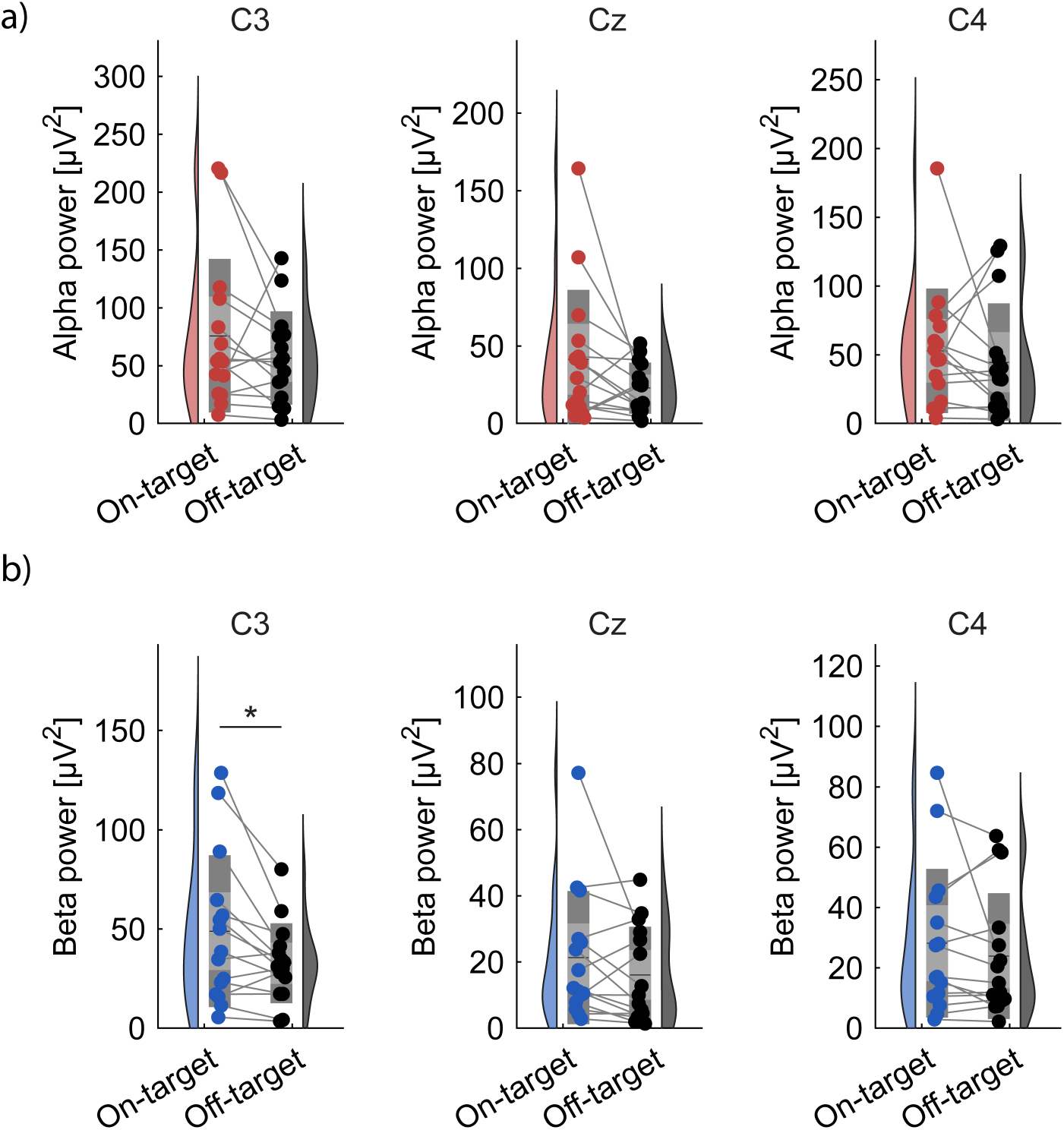
Alpha- and beta-band power differences between successful and unsuccessful kicks. (a) Participant-level alpha-band power (8–15 Hz) after pooling trials across recording days. No alpha-band comparison survived Holm correction (C3: Holm-adjusted p = 0.381; Cz: Holm-adjusted p = 0.191; C4: Holm-adjusted p = 0.528). (b) Participant-level beta-band power (15–30 Hz). Beta-band power was significantly greater before successful than unsuccessful kicks only at C3 after Holm correction (C3: two-sided p = 0.0139, Holm-adjusted p = 0.0418), whereas Cz and C4 were not significant (Cz: Holm-adjusted p = 0.375; C4: Holm-adjusted p = 0.524).

## Discussion

This study demonstrates that pre-movement EEG activity recorded with a low-density, portable three-channel system contains information related to the subsequent outcome of futsal free kicks.

Within-day analyses showed that the linear SVM classifier decoded successful and failed kicks above the corresponding label-shuffled null distributions across C3, Cz, and C4. In contrast, the same decoding approach did not generalize reliably across recording days, indicating that the discriminative patterns were not stable across sessions. A complementary band-power analysis further identified an increase in C3 beta-band power before successful kicks. Together, these findings suggest that successful and failed kicks are preceded by reproducible within-day differences in pre-movement sensor-level EEG activity, while also highlighting the session-specific nature of these neural signatures.

The present findings are consistent with a broader literature showing that brain activity during the preparatory period is related to subsequent skilled motor performance. EEG studies of skilled psychomotor performance have long suggested that successful performance is associated with distinct pre-action neural states, particularly in tasks requiring precise timing, attention, and motor control[18]. In sport-specific contexts, pre-shot or pre-movement EEG rhythms have been associated with performance outcomes in precision sports such as shooting and golf putting[12,14]. The present study extends this line of work by examining a more dynamic, whole-body sports action performed in a gymnasium setting rather than a tightly constrained laboratory task. Thus, the current results support the view that the neural state immediately before movement is not merely a passive correlate of performance, but a measurable component of the motor-preparatory process that can help distinguish successful from unsuccessful actions.

A key contribution of this study is that outcome-related neural information was detected using only three central EEG channels. Many EEG studies of motor performance have relied on high-density electrode montages or highly controlled laboratory environments, which can be difficult to translate into routine sports practice. Practical sports applications require systems that can be set up quickly, tolerate natural movement, and be repeatedly used during training without imposing excessive burden on athletes. This motivation is consistent with recent arguments that dry or low-burden EEG systems may help move sport neuroscience from laboratory settings toward field-relevant monitoring[19]. By showing that pre-movement EEG from C3, Cz, and C4 contained decodable information and that C3 beta-band power showed an outcome-related spectral difference, this study provides an initial step toward field-oriented brain-state monitoring in sport[20].

The within-day decoding results provide evidence that single-trial pre-movement EEG contains outcome-related information even when recorded with a minimal three-channel montage during a field-relevant whole-body sports action. The SVM classifier exceeded the shuffled-label null distributions across all three central channels, indicating that the effect was not confined to a single electrode or an isolated analysis choice. This is notable because EEG classification remains challenging even in standard brain-computer interface settings, where signal-to-noise ratio, inter-trial variability, and limited training data often constrain classifier performance[21]. Although the absolute AUC values were modest and are not yet sufficient for stand-alone practical prediction, their consistent elevation above participant- and session-specific shuffled-label distributions across all three channels indicates that the recorded signals contained detectable information related to subsequent kick outcome.

Although the behavioral task involved a lower-limb action, successful decoding from C3, Cz, and C4 is physiologically plausible because kicking is not an isolated movement of the leg. Soccer and futsal kicking require coordinated control of the trunk, pelvis, support leg, kicking leg, and upper limbs to regulate balance, approach, impact mechanics, and movement direction[22]. Therefore, EEG signals recorded over central sensorimotor regions may reflect the integrated motor-preparatory state for the whole action rather than activity related to a single effector. At the same time, because the present study used a sparse scalp montage, the observed channel-level effects should not be interpreted as precise source-localized activity. Instead, C3, Cz, and C4 should be treated as practical sensor-level markers that capture information relevant to whole-body motor preparation.

The failure of the SVM model to generalize across days is an important result for practical deployment. The within-day results indicate that outcome-related information was present in the EEG signals, whereas the cross-day results show that this information was not represented in a session-invariant form. This pattern is consistent with a major challenge in EEG-based decoding: signal distributions can change across sessions because of differences in electrode placement, impedance, fatigue, attention, arousal, movement artifacts, and environmental conditions. In brain-computer interface research, such non-stationarity has motivated transfer learning, adaptive normalization, and domain-adaptation approaches to reduce calibration burden across subjects and sessions[23,24]. Therefore, the limited cross-day generalization indicates that realistic sports EEG systems will likely require day-specific calibration or adaptive decoding. In near-term applications, an individualized within-session or short-calibration workflow may be more realistic than the immediate use of a fixed model across recording days.

The C3 beta-band increase before successful kicks provides a physiologically interpretable complement to the decoding results. Event-related changes in sensorimotor rhythms are a fundamental feature of EEG during motor preparation and execution[25]. In particular, beta-band oscillations have been linked to the maintenance of the current sensorimotor or cognitive set[26]. In the present task, higher C3 beta-band power before successful kicks may reflect a relatively stable preparatory state, in which the intended action plan was maintained before movement execution. The channel- and frequency-specific nature of this effect argues against a purely global change in signal amplitude. However, beta activity is context dependent and can vary with task demands, learning state, and adaptation requirements[27]. Thus, the present C3 beta-band finding should be interpreted as an outcome-related sensor-level correlate of successful preparation in this futsal kicking task, rather than as a causal determinant of success.

The decoding and band-power analyses provide complementary views of the same underlying problem. The SVM analysis tested whether the pre-movement EEG waveform contained information sufficient to discriminate successful and failed kicks at the single-trial level. In contrast, the band-power analysis tested whether more interpretable spectral features, including alpha- and beta-band power, differed between outcome conditions at the participant level. The combination of these two analyses is useful because a waveform-based classifier can indicate that discriminative information is present, whereas interpretable oscillatory features can suggest what aspect of the pre-movement state might be monitored in future applications. This distinction is especially relevant for neurofeedback and closed-loop training, where a target feature should be measurable, interpretable, and behaviorally meaningful[28,29]. Previous work suggests that sensorimotor rhythm neurofeedback can influence motor behavior and sport performance, although target selection and intervention design remain important challenges[10,30]. The present findings therefore suggest a possible future workflow in which a short calibration period is used to estimate an athlete’s day-specific preparatory EEG state, after which classifier outputs or C3 beta-band power could be monitored during practice and potentially used for feedback or cueing. However, the present study did not test an intervention, and whether modifying these EEG states can improve sport-specific performance remains to be evaluated directly.

Several limitations should be considered. First, although the sample size was modest, it was comparable to or larger than the sample sizes reported in previous repeated-measures sport-EEG studies[12,14]. Moreover, each participant completed 100 trials across two recording days. The sample was limited to young right-footed male participants with soccer experience; therefore, generalization to female athletes, left-footed players, elite players, or novices remains unknown. The study was designed to compare successful and unsuccessful trials within the same soccer-experienced participants, rather than to compare different levels of expertise. Accordingly, the absence of a novice or non-soccer control group limits conclusions about expertise specificity, but does not affect the primary within-participant comparison. Second, the task involved a fixed target and did not reproduce the full perceptual and tactical demands of actual gameplay. However, many passes in actual soccer and futsal matches are performed without a multi-step run-up, and the single-step approach therefore retained a match-relevant short-approach kicking pattern. The present findings are most directly applicable to such short-approach passes and kicks rather than to long-run-up shots. Third, the three-channel EEG montage improved practicality but limited spatial resolution and prevented source localization. Fourth, movement onset was defined from head acceleration, the analysis was restricted to the preceding 2-s interval, and waveform-based quality checks were performed. Nevertheless, because simultaneous electrooculography (EOG) and electromyography (EMG) were not recorded, subtle movement-related, ocular, or muscular contributions cannot be completely excluded. The findings should therefore be interpreted as sensor-level EEG associations rather than as source-localized cortical effects. Fifth, the cross-day results show that the present models are not yet robust enough for calibration-free longitudinal use. Finally, the beta-band effect is correlational; it shows an association between C3 beta-band power and successful kicks but does not prove that changing this activity would improve performance. Future studies should test larger and more diverse athlete populations, incorporate multimodal recordings such as EMG, motion capture, video, and head acceleration controls, evaluate whether movement-only signals can explain any part of the decoding performance, and develop adaptive decoding methods that can handle session-to-session variability. Intervention studies will also be needed to determine whether feedback or closed-loop cueing based on pre-movement EEG features can improve sport-specific performance.

## Acknowledgments

The authors thank Shoko Tonomoto, Aya Kamiya, and Sayoko Ishii for their assistance.

## Author contributions

Conceptualization: S.H.

Methodology: S.H., S.I.

Investigation: S.H.

Data curation: S.H.

Formal analysis: S.H.

Visualization: S.H., S.I.

Writing – original draft: S.H.

Writing – review and editing: S.I.

Supervision: S.I, J.U.

Funding acquisition: S.I., J.U.

## Data availability

The individual-level EEG and behavioral data underlying the findings of this study cannot be made publicly available because the informed consent obtained from the participants did not authorize unrestricted public sharing of these data. De-identified data may be made available to qualified researchers upon approval by the Ethics Committee of the Faculty of Science and Technology, Keio University, and completion of an appropriate data use agreement. Data access requests should be directed to the Ethics Committee Office, Faculty of Science and Technology, Keio University, c/o the Environmental Conservation Center Office (; telephone: +81-45-566-1455; postal address: Room 202, Building 24, 3-14-1 Hiyoshi, Kohoku-ku, Yokohama 223-8522, Japan). Requests should reference ethics approval numbers 2023-141 and 2024-011.

All author-generated analysis code underlying the findings of this study is publicly available at https://github.com/HiroseSadao/futsal-eeg-analysis under the MIT license.

